# Evolutionary engineering a larger porin using a loop-to-hairpin mechanism

**DOI:** 10.1101/2023.06.14.544993

**Authors:** Rik Dhar, Alexander M. Bowman, Brunojoel Hatungimana, Joanna SG Slusky

## Abstract

In protein evolution, diversification is generally driven by genetic duplication. The hallmarks of this mechanism are visible in the repeating topology of various proteins. In outer membrane β-barrels, duplication is visible with β-hairpins as the repeating unit of the barrel. In contrast to the overall use of duplication in diversification, a computational study hypothesized evolutionary mechanisms other than hairpin duplications leading to increases in the number of strands in outer membrane β-barrels. Specifically, the topology of some 16- and 18-stranded β-barrels appear to have evolved through a loop to β-hairpin transition. Here we test this novel evolutionary mechanism by creating a chimeric protein from an 18-stranded β-barrel and an evolutionarily related 16-stranded β-barrel. The chimeric combination of the two was created by replacing loop L3 of the 16-stranded barrel with the sequentially matched transmembrane β-hairpin region of the 18-stranded barrel. We find the resulting chimeric protein is stable and has characteristics of increased strand number. This study provides the first experimental evidence supporting the evolution through a loop to β-hairpin transition.

**Highlights:**

- We find evidence supporting a novel diversification mechanism in membrane β-barrels
- The mechanism is the conversion of an extracellular loop to transmembrane β-hairpin
- A chimeric protein modeling this mechanism folds stably in the membrane
- The chimera has more β-structure and a larger pore, consistent with a loop-to-hairpin transition

## Introduction

Outer membrane proteins (OMPs) are found in the outer membrane of Gram-negative bacteria, mitochondria, and the chloroplast [1, 2]. Unlike inner membrane proteins which consist of membrane-spanning alpha-helices, OMPs generally consist of membrane-spanning anti-parallel β-barrels, connected sequentially with shorter turns in the inter-membrane side and longer loops on the outward side [3–5]. The loops of OMPs have various functions including structural interaction with the lipopolysaccharides [6], channel gating [7, 8], protease activity [9], interaction with target protein [10], and inter-barrel interactions [11].

OMPs differ from each other by the number of strands forming their transmembrane β-barrel domain, which ranges from 8 to 36 strands [12, 13]. Similar to divergent protein evolution more generally [14–22], OMPs primarily diversified through genetic duplication of a primordial peptide specifically an ancestral β-hairpin peptide [23]. As a result, the β-hairpin motif forms the repeating unit in OMPs, which is evident from internal sequence similarities [24–26]. Analysis of these internal sequence repeats in structurally resolved OMPs revealed an overview of the OMP evolutionary pathway of OMPs tracing the relationship between β-barrels classes of different strand numbers [21].

In addition to duplication events, further diversification occurred by mutation, deletion, and insertion followed by further gene duplication [20, 27] creating a relationship web among most outer membrane proteins. Among the most thought-provoking of these relationships was that transmembrane strands are not only related to each other, but some are also related to extracellular loops. The presence of sequence alignment between β-hairpins of 18-stranded β-barrels and the loops of 16-stranded β-barrels led to the hypothesis that extracellular loops are sometimes evolutionarily converted into transmembrane β-hairpins. This mechanism might be particularly surprising because it is a non-duplication event causing a repeat protein topology. Moreover, soluble and transmembrane regions are not generally thought to be interchangeable. However, since outer membrane proteins and soluble proteins have similar hydrophobicities and are just “inside out” [28], this hypothesized mechanism may in fact occur in nature.

Due to the importance of the duplication evolutionary mechanism in OMP diversification, many have worked to engineer OMPs using this principle [19, 29–32]. However, incorporating additional strands into β-barrels is challenging, as evidenced by the unsuccessful integration of additional strands in OmpG [33] and the subsequent loss of channel conductance in LamB after the insertion of additional β-hairpin [32]. Another laboratory approach for strand accretion involves creating chimeras from two different β-barrels [19, 32, 34, 35]. Again this is also challenging as not all combinations result in a stable protein [35, 36] nor do they result in oligomeric proteins [19, 34]. The interaction of neighboring residues [37], the sensitivity of residues to mutations [38] and the role of hydrogen bonding in the backbone [3], can frustrate design attempts involving strand insertion.

OMP engineering is most likely to be successful when natively adjacent strands are inserted, such as the first and the last strand, in 8 to 16-stranded OmpX engineering [19] or when strands are replaced with corresponding ones in the OmpF and OmpG chimera [34]. These approaches preserve the interaction between conserved residues, ensuring the stability of the resulting protein. Because of this, usage of evolutionary relationships is known to be more likely to yield success in the difficult endeavor of enlarging OMP β-barrels.

Here we use the hypothesized loop-to-hairpin mode of evolution to engineer an 18-stranded β-barrel from a 16-stranded β-barrel. We selected a protein pair from our previous study that shows evidence for strand accretion through loop-to-hairpin transition [21]. This protein pair consists of the 16-stranded β-barrel PorB from *Neisseria Gonorrhoeae* (PDBID 4AUI) [39]. and 18-stranded β-barrel major outer membrane protein (MOMP) from *Campylobacter jejuni* (PDBID 5LDV) [40]. Loop L3 of PorB is its longest loop which lines the lumen of the barrel domain [39]. This loop was found to align with a β-hairpin of MOMP. We engineered a chimeric protein by replacing loop L3 of PorB with the aligned region of MOMP. Despite being composed of proteins from different species with only 21.7% sequence identity, the chimeric protein stably expresses, folds, and inserts into the outer membrane. Using secondary structure estimation from circular dichroism and pore size estimates from single channel conductance, the chimeric protein most likely forms an 18-stranded barrel.

## Results

### Chimeric protein folds in the outer membrane

To engineer an 18-stranded barrel from a 16-stranded barrel based on loop-to-hairpin evolution we created a chimeric protein from PorB and MOMP. This chimeric protein, which we call PorB_18_, was created by replacing the loop L3 of wild-type PorB (PorB_WT_) with the sequentially aligned region of MOMP (Fig. 1A–C). The insert from MOMP in chimeric protein PorB_18_ that replaces the long loop L3 of PorB_WT_ consists of the MOMP short loop L3, the two MOMP strands S7 and S8, the MOMP long loop L4, and the MOMP turn T3 (Fig. 1C). To evaluate the stability of the chimeric protein, we tested its production in the outer membrane. For this, the N-terminal signal sequence from the *Escherichia coli* OMP OmpA was added to the N-terminal of PorB_WT_ and PorB_18_ to target the proteins to the outer membrane [41, 42]. For expression, we used an *E. coli* strain that lacks the periplasmic protease DegP, as neisserial porins are difficult to express recombinantly in the *E. coli* outer membrane [43, 44]. We lysogenized this ΔDegP strain with λDE3 which allows for the use of the pET expression system for the ‘leaky’ expression of PorB_WT_ and PorB_18_ [45].

**Figure 1:**
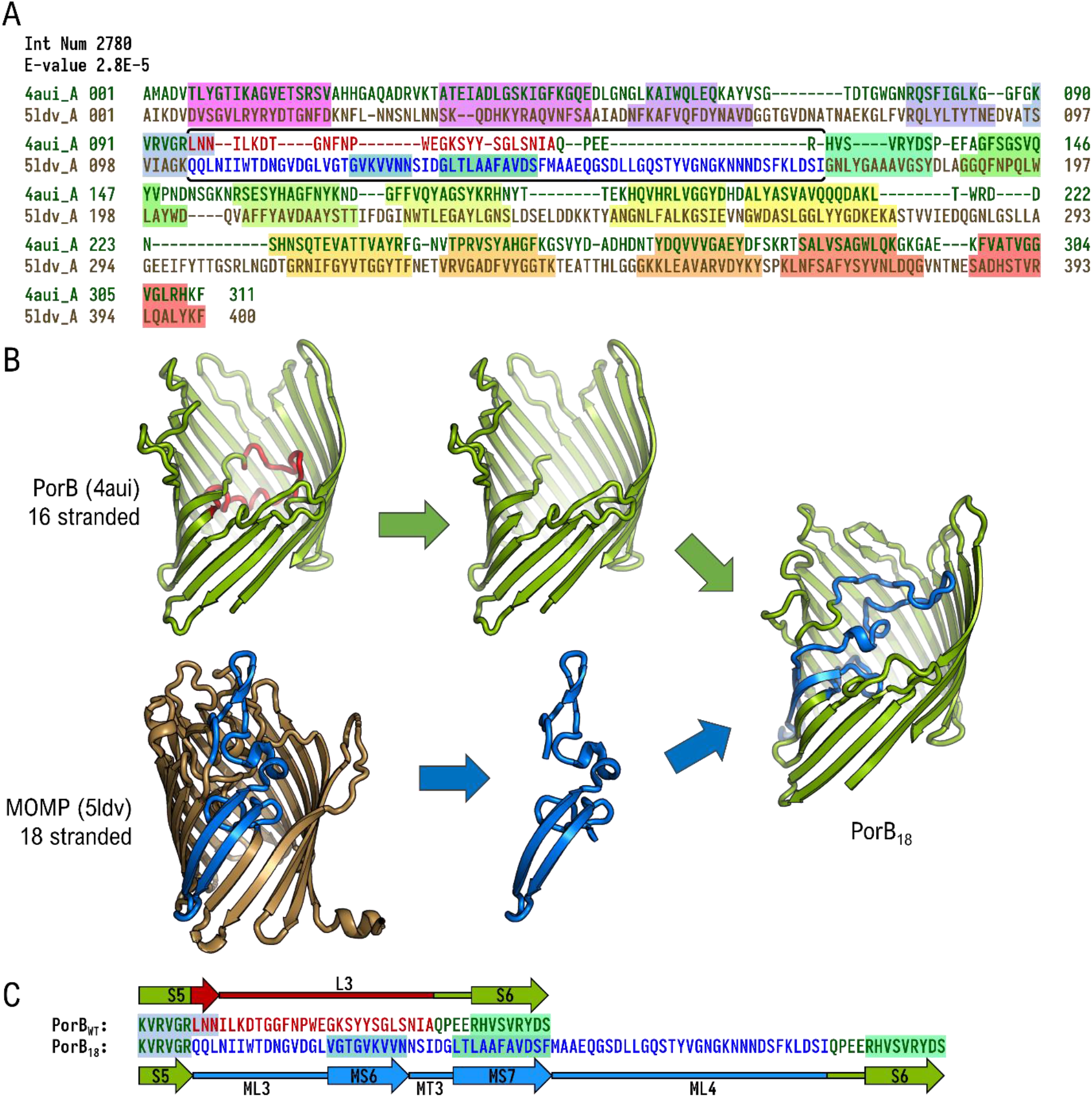
PorB_18_, a chimera of PorB_WT_ and MOMP: **(A)** Sequence alignment between 16-stranded PorB (PDB ID 4aui [39], green text) and 18-stranded MOMP (PDB ID 5ldv [46], brown text). (Adapted from Franklin [47]) The highlighted colors represent the β-strands as identified from the crystal structures: the N-terminal strand is in magenta, the C-terminal strand is in red, and the intermediate strands are highlighted in a rainbow gradient. The horizontal double bracket indicates the region of the loop/hairpin alignment **(B)** Construction of the chimeric protein PorB_18_. The majority of loop L3 (red) from PorB_WT_ (green) has been removed, and the aligned peptide sequence (blue) of MOMP has been inserted (brown) in L3’s position. **(C)** Sequence comparison between PorB_WT_ and PorB_18_ showing the modified region. The deleted region is shown in red and the inserted MOMP region is shown in blue.

Western blot of different subcellular fractions of *E. coli* showed that PorB_WT_ and PorB_18_ are expressed and are trafficked to the outer membrane (Fig. 2A). To assess the proper folding and insertion of the recombinant proteins in the outer membrane, we carried out proteinase K digestion in bacterial cells. Proteinase K is a serine protease that predominantly cleaves the peptide bond adjacent to the carboxylic group of an aliphatic or aromatic residue [48]. Proteinase K assay has been used in the study of membrane proteins to discover exposed or disordered regions [49] and to evaluate OMP insertion in the outer membrane [50]. *E. coli* expressing the recombinant proteins were permeabilized and treated with proteinase K for 20 minutes on ice. Western blot of these proteinase K treated cells indicated both PorB_WT_ and PorB_18_ were resistant to proteinase K digestion (Fig. 2B) as indicated by the small amount of band density reduction after digestion. We used TolC as a control for this assay as it has a 5 kDa disordered region in the periplasm that is susceptible to proteinase K digestion [51]. PorB_WT_, PorB_18_, and TolC were found to have a low level of degradation products, which can be attributed to the endogenously expressed OmpT protease in the *E. coli* strain used for expression. Resistance to proteinase K digestion indicates that both PorB_WT_ and chimeric PorB_18_ insert and fold into the outer membrane.

**Figure 2:**
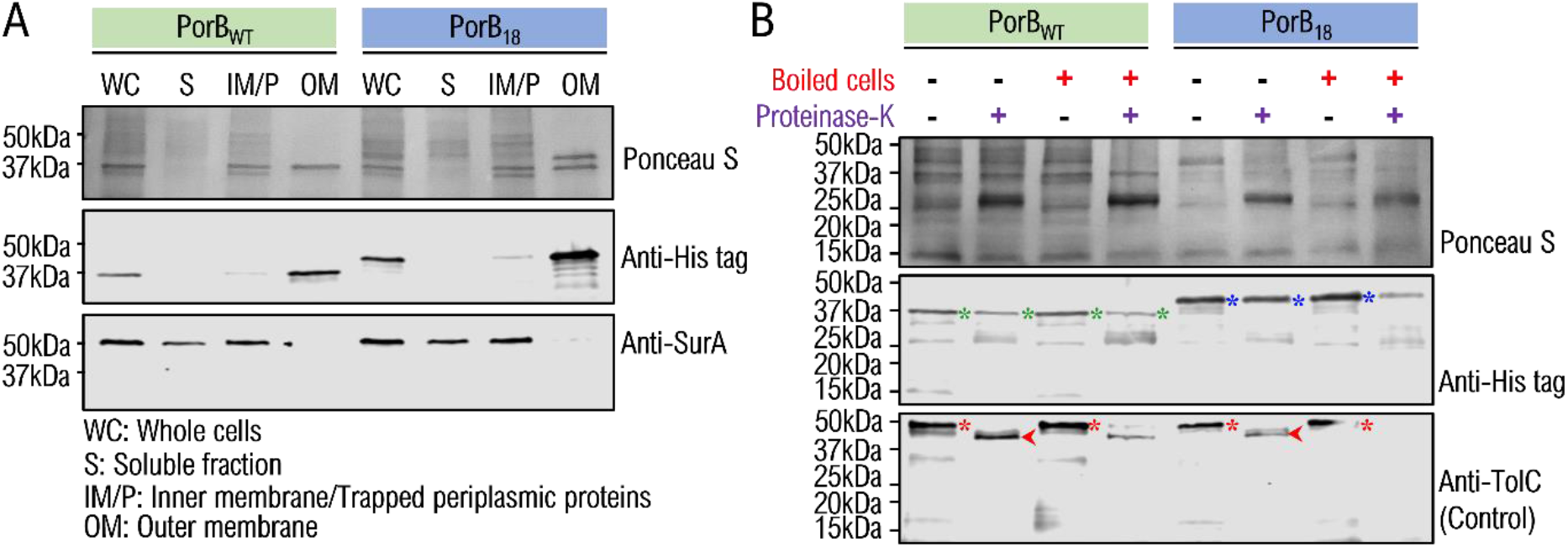
Expression of PorB_18_ in *E. coli* outer membrane: **(A, B)** Western blots with the nitrocellulose membrane stained with Ponceau S and subsequently immunoblotted against various antibodies. **(A)** Western blot of different fractions of *E. coli* displaying the localization of the his-tagged recombinant proteins PorB_WT_ and PorB_18_ in the outer membrane. SurA, a periplasmic protein, indicates the localization of periplasmic proteins. Whole-cell fractions (WC), soluble (S), inner membrane and trapped periplasmic protein (IM/P), and outer membrane (OM) **(B)** Western blot displaying the susceptibility of proteins to proteinase K under different conditions. His-tagged PorB_WT_ (blue asterisks) and PorB_18_ (green asterisks) show resistance to proteinase K when not boiled but are partially susceptible when heat denatured. TolC (red asterisks) is used as a positive control as it shows a 5 kDa cleavage in the presence of proteinase K resulting in a 45 kDa cleavage product (red arrows) [51].

### Chimeric protein has increased β-sheet content

To estimate the secondary structure of the chimeric protein PorB_18_ we measured and analyzed the circular dichroism (CD) spectra of these proteins. Because a higher concentration of protein was required for CD analysis, the recombinant proteins were expressed as inclusion bodies, purified, and refolded into lauryldimethylamine oxide (LDAO) detergent micelles [52]. Expression as inclusion bodies provides a much higher yield compared to expression in the outer membrane. The refolded proteins were separated by size exclusion chromatography (SEC) to isolate the folded trimeric fractions. We treated the eluents with a reversible crosslinker dithiobis succinimidyl propionate (DSP) (Fig. 3A) to verify the quaternary structure. Crosslinking and subsequent visualization by SDS-PAGE indicated that both the recombinant proteins PorB_WT_ and PorB_18_ trimerized (Fig. 3B). It is therefore likely that the inserted region from MOMP in the chimeric protein PorB_18_ does not destabilize the quaternary structure.

**Figure 3:**
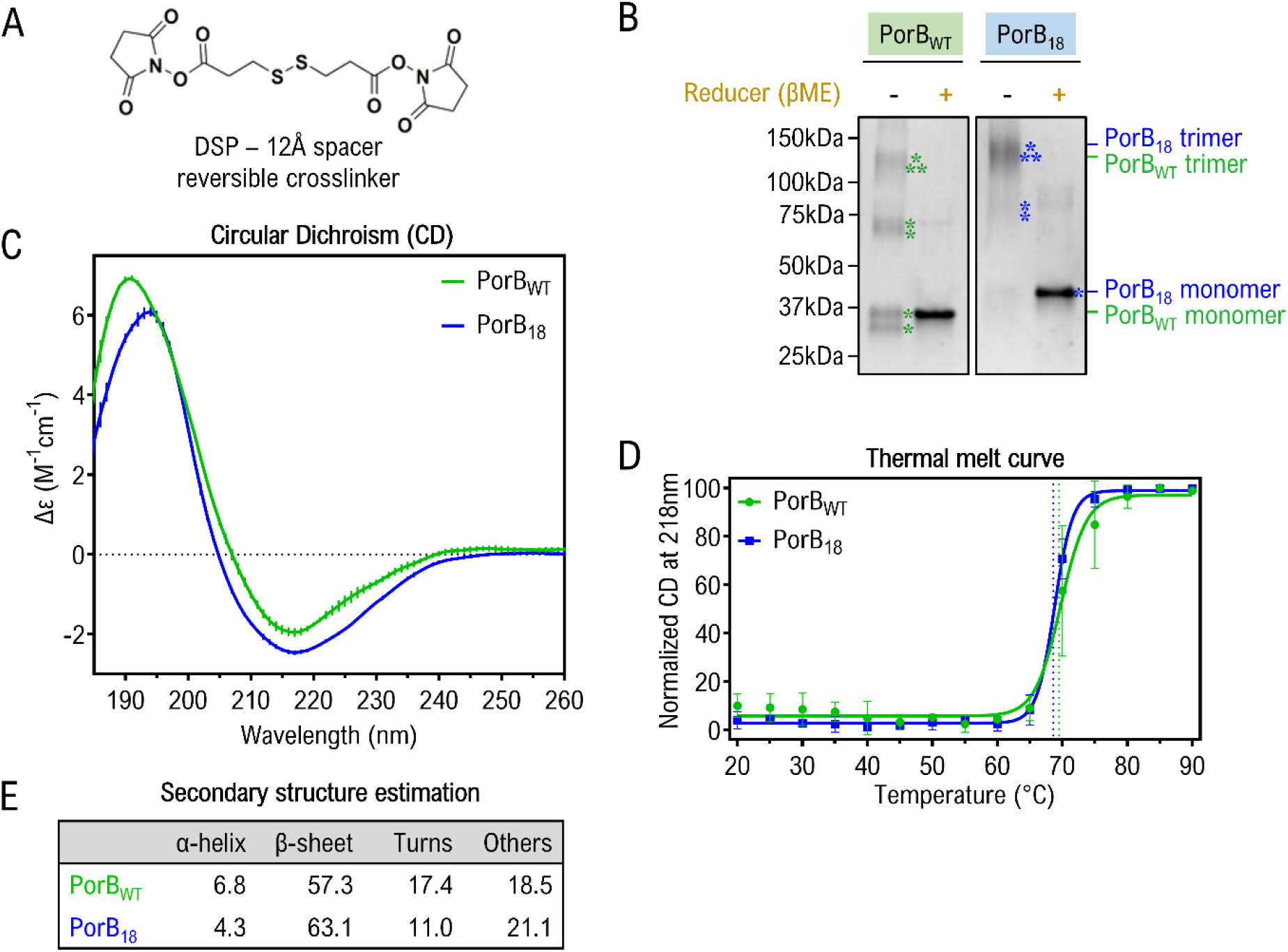
Secondary and tertiary structural analysis of PorB_WT_ and PorB_18_: **(A)** Reversible crosslinker DSP. **(B)** SDS-PAGE displaying DSP crosslinked purified proteins, with samples either untreated or treated with the reducer, β-mercaptoethanol to reverse crosslinking. Bands corresponding to PorB_WT_ (green) and PorB_18_ (blue). monomers are marked with (⁎), dimer with (⁑), and trimer with (⁂). **(C)** CD spectra of the proteins with error bars indicated by vertical lines. **(D)** Normalized CD spectra at 218 nm show the thermal unfolding of PorB_WT_ (green) and PorB_18_ (blue). The lines represent the regression curve with a sigmoidal fit. Dotted vertical lines indicate the T_m_. **(E)** Secondary structure estimation from the CD spectra.

CD spectra of PorB_WT_ and PorB_18_ revealed both proteins have β-sheet-like spectra with minima at 218 nm (Fig. 3C). The PorB_18_ spectra had higher β-sheet content than the PorB_WT_ as determined by the JASCO multivariate SSE program (Fig. 3E). This supports that the inserted region is adding extra β-strands to the structure. The proteins exhibited similar thermal stability (T_m_ for PorB_WT_ = 69 °C, T_m_ for PorB_18_ = 68 °C) as determined by tracking temperature-dependent unfolding through CD absorption measurement at 218 nm (Fig.3D).

### Chimeric β-strands are buried

To assess if the extra β-strands are exposed or buried, PorB_WT_ and PorB_18_ folded in detergent micelles were subjected to proteinase K for a longer duration. Proteins in detergent micelle are likely to be more susceptible to proteinase K digestion as they are more exposed than in the outer membrane, where the LPS shields them. We found PorB_18_ to be more susceptible to proteinase K than PorB_WT_ (Fig. 4A – B) as it is fully cleaved after 60 minutes and PorB_WT_ is not. To evaluate which regions of the proteins are exposed and are susceptible to proteinase K, we analyzed the cleavage products with matrix-assisted laser desorption/ionization-time of flight (MALDI-TOF) mass spectrometry (Fig. 4C & D).

**Figure 4:**
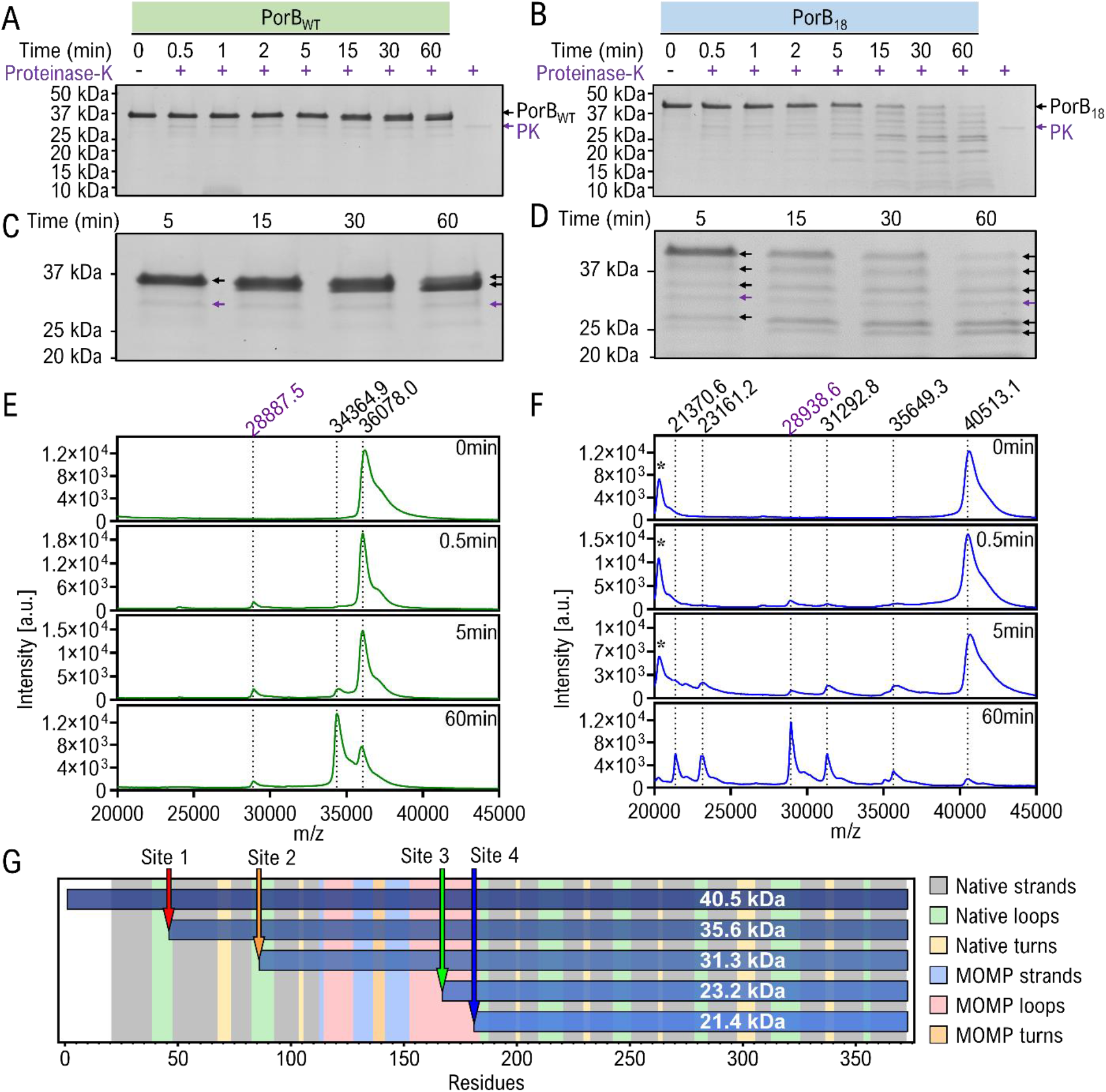
Proteinase K digestion of PorB_18_ refolded in micelles. SDS-PAGE of **(A)** PorB_WT_ and **(B)** PorB_18_ refolded in micelles, treated with proteinase K for different durations. An enlarged view of the lanes from 5 min to 60 min is shown in panel **(C)** for PorB_WT_ and **(D)** for PorB_18_. The black arrow marks the band that appears as a peak in MALDI-TOF. The purple arrow marks the proteinase K band. MALDI-TOF of **(E)** PorB_WT_ (green) and **(F)** PorB_18_ (blue) treated with proteinase K for different durations. The numbers above the graph are the mass-by-charge ratio of the peaks. Peaks indicated by black asterisks are doubly charged peaks of full-length PorB_18_ seen at 40513.1. Peaks indicated by purple numbers are proteinase K which has a molecular weight of 28.9 kDa. **(G)** Mass spectrometry-based predicted proteinase K sites for PorB_18_. The horizontal bars represent the predicted proteinase K digestion product for each mass spectrometry peak from Figure 4. The x-axis shows the residue numbers and the left and right edges of the bars represent the N- and C-terminal residues. The colored vertical region represents the secondary structures

Analysis of the cleavage products by MALDI-TOF mass spectrometry indicated one major cleavage product for PorB_WT_ (Fig. 4E) at 34,364.9 Da. This product is difficult to differentiate from the uncleaved protein band on SDS-PAGE gel at this figure size but can be more clearly seen in a larger version (Fig. 4C). All possible proteinase K cleavage sites were predicted using Expasy PeptideCutter [53] for PorB_WT_ and PorB_18_ (Fig. S1A and B). Using the predicted cleavage sites and the peaks from the mass spectra, the proteinase K cleavage site for PorB_WT_ is identified as residue L16 on the N-terminal tag (Fig. S1C). The peak at 28,887.5 Da is likely proteinase K with a known molecular weight of 28.9 kDa.

In contrast to the single cleavage product from PorB_WT_, four major cleavage products are formed for PorB_18_. (Fig. 4F). The additional peak at 20.2 kDa evident in 0 – 5 minute time points is the doubly charged uncleaved protein. While PorB_18_ is protected against proteinase K in bacterial membranes (Fig. 2B), the increased susceptibility to proteinase K in micelles is likely due to the inserted loop region having exposed hydrophobic residues in the detergent micelles. These residues which are likely to be shielded in the outer membrane by LPS are exposed in the detergent micelles.

To identify the cleavage sites in PorB_18_ that result in the four major cleavage products, MALDI-TOF spectra of the 60 min sample was acquired at 5k to 45k mass range (Fig. S2A). Three smaller size cleavage products were identified of mass 12.6 kDa, 9.9 kDa, and 8.3 kDa that correspond to the mass differences between the four major cleavage products. Two of the smaller cleavage products (8.3 kDa and 12.6 kDa) showed similar MALDI-TOF spectral pattern (Fig. S2B and C) indicating that both these cleavage products end with the amino acids serine and threonine. The possible cleavage sites (Fig. S1B) were refined by using the information from the extended MALDI-TOF spectra (Fig. S2A – C) to determine the sequences for the four major cleavage products (Fig. S2D and 4G).

Despite having multiple proteinase K cleavage sites (Fig. S1B), none of the calculated products start or end in the MOMP β-strands region (Fig. 4G & S2E). All cleavage sites are in the extracellular loop region and would be likely to be shielded by the extensive sugars of LPS in cells and therefore uncleaved as previously shown (Fig. 2B). However, micelles and the lipids would not shield these sites and they would therefore subject to cleavage as previously shown (Fig. 4D). These positions are also all in loop regions which tend to be more flexible and therefore more likely to become exposed to proteinase. Though the two smaller proposed cleavage products are a result of cleavage in the added region of PorB18, they are in the loops of those regions, not in the strands of those regions. Therefore, the inserted β-strands are likely to be buried in the structure.

### 18-stranded predicted structure agrees with experimental observations

Although the CD spectra show that the inserted region in PorB_18_ is forming β-strands and the MALDI-TOF analysis of proteinase K indicates that those strands are buried in the structure, neither demonstrates that these extra β-strands are becoming part of the β-barrel. Through ColabFold [54] we used AlphaFold-multimer [55, 56] to predict the structure of the chimeric protein (Fig. 5A and B). The top two models both predict the inserted region to be buried within the structure, but one predicts the entire inserted region along with the β-strands to be inside the lumen of the β-barrel forming a plugged 16-stranded β-barrel (Fig. 5A) while the second predicts the inserted β-strands to be part of the β-barrel forming an 18-stranded β-barrel with a long loop lining the lumen of the barrel (Fig. 5B). Notably, the confidence of the prediction as measured by the predicted local distance difference test (pLDDT) is very low for the inserted region for both predictions. However, the prediction confidence was higher when it predicted the structure to be 18-stranded (Fig. 5C).

**Figure 5:**
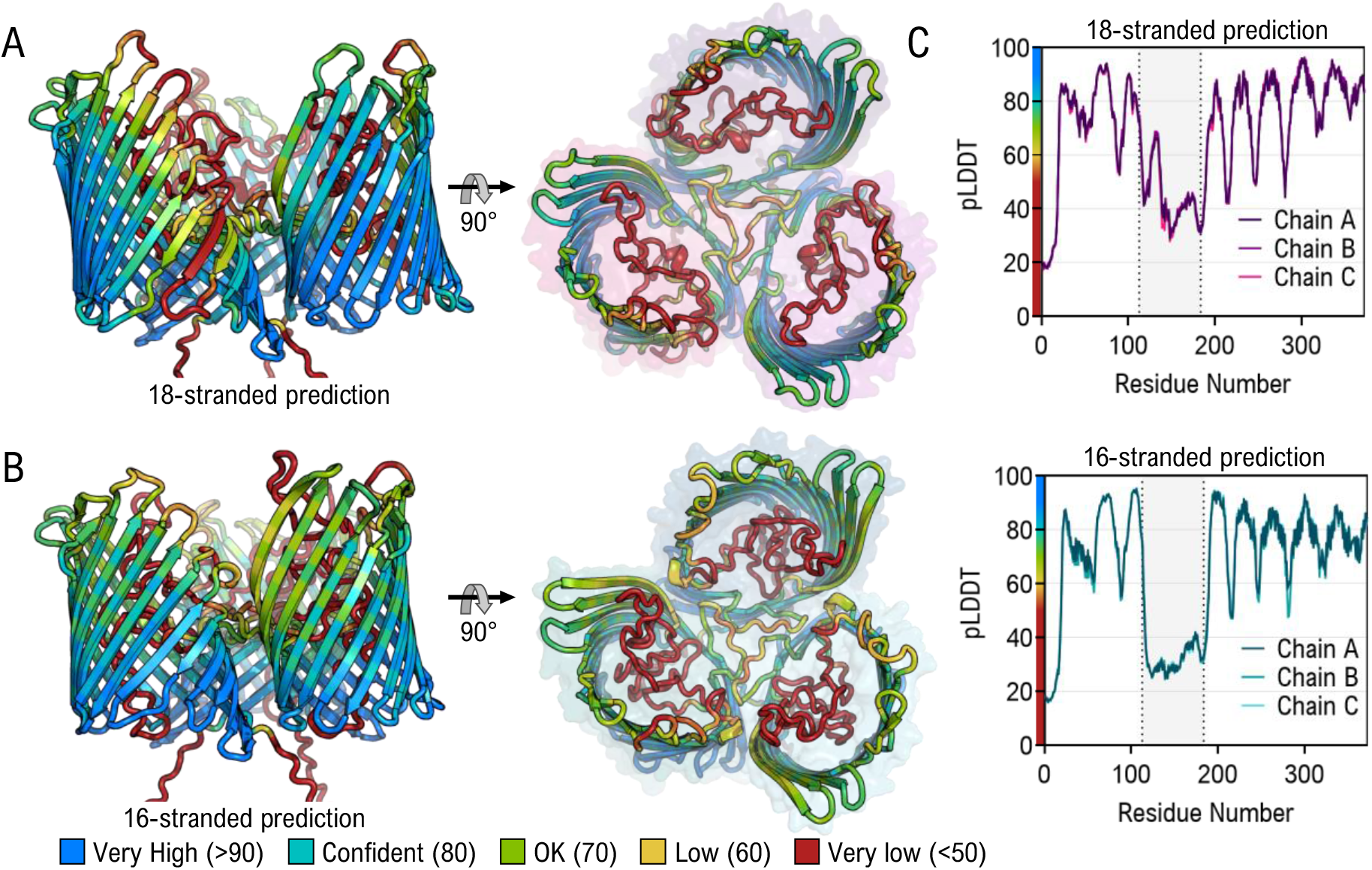
AlphaFold2 prediction of PorB_18_. **(A and B)** Top two ranked predicted structures of PorB_18_: **(A)** 18-stranded prediction of PorB_18_, **(B)** 16-stranded prediction of PorB_18_. **(C)** Plots showing pLDDT scores per residue for each of the two predicted structures. The shaded grey area shows the insert region.

To further verify which of the predicted structures is more likely, we analyzed the predicted structures to see which of the two structures agrees more with the experimental data. First, we mapped the proteinase K cleavage sites predicted through mass spectrometry (Fig. 4G) onto the two predicted structures (Fig. 6A and B). While cleavage site 1 is exposed in both PorB_18_ predicted structures compared to PorB_WT_ (Fig. 6C), site 2 is exposed only in the 18-stranded prediction of PorB_18_. Next, the theoretical CD spectra for the AlphaFold models were estimated using the PDB2CD web server [57] (Fig. 6D). The 18-stranded PorB_18_ model was estimated to be 61.7% β-sheet, and the 16-stranded model was estimated to be 57.6% β-sheet. The theoretical CD for the PDB structure of PorB_WT_ (PDB ID 4aui) was calculated to be 55.0% β-sheet. Based on the mapped proteinase K sites and the theoretical CD spectra, the 18-stranded predicted structure of PorB_18_ agrees more with observed experimental data.

**Figure 6:**
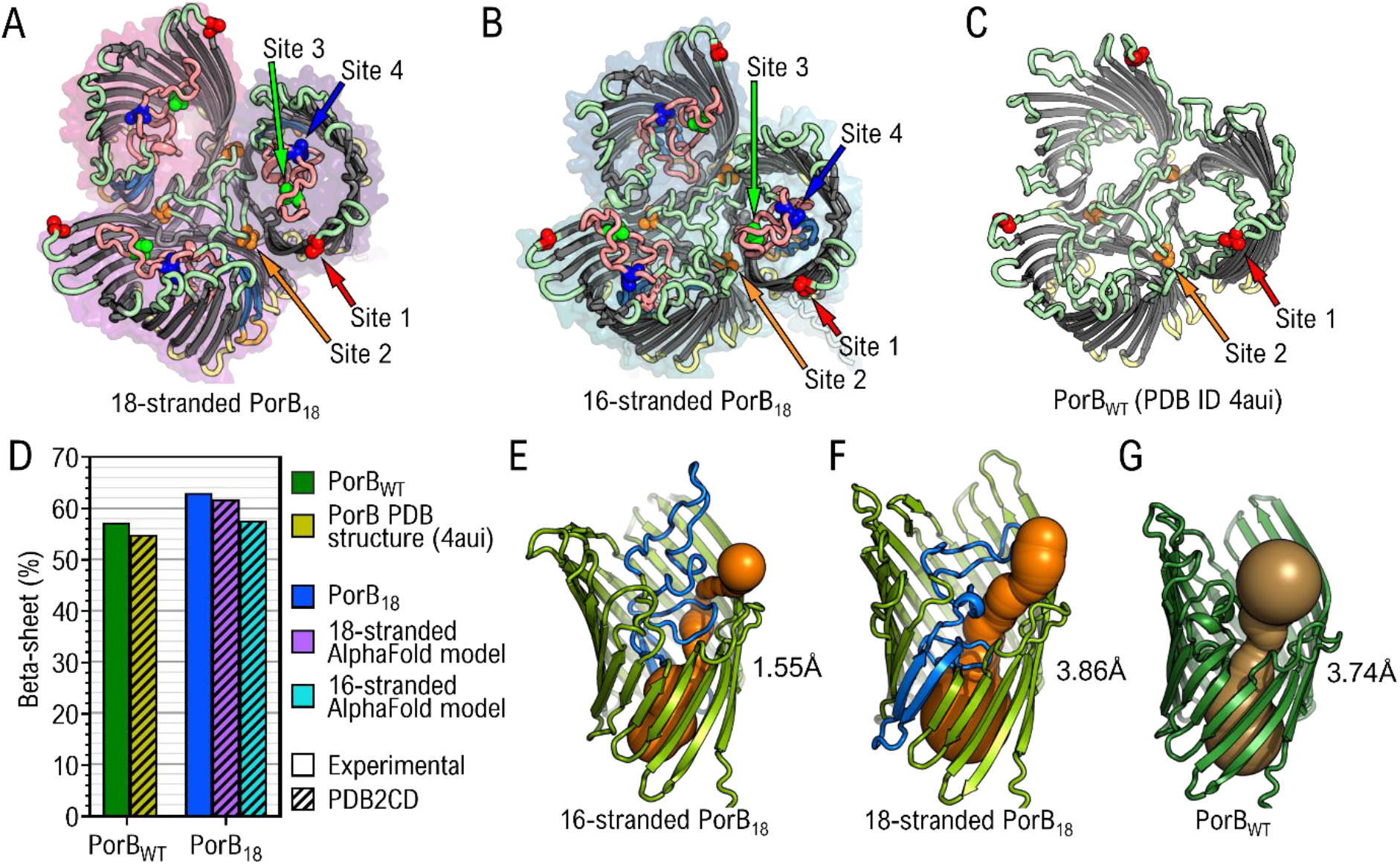
Analysis of predicted structures of PorB_18_. **(A – C)** Residues with the predicted proteinase K cleavage sites are shown as spheres in **(A)** 18-stranded prediction of PorB_18_, **(B)** 16-stranded prediction of PorB_18_, and **(C)** PorB_WT_ (PDB ID 4aui) **(D)** The predicted β-sheet percentage calculated from the experimental CD data of PorBWT and PorB18 is compared with the predicted β-sheet percentage from theoretical CD spectra of the PDB structure of PorBWT and the AlphaFold models calculated using the PDB2CD server [57]. **(E – G)** Channel prediction using CAVER 3.03 for 16-stranded PorB_18_ **(E)**, 18-stranded PorB_18_ **(F)**, and PorB_WT_ **(G)**. The number next to the cartoon is the bottleneck radius.

### Chimeric strands increase the pore radius

To investigate the pore size of the chimeric protein PorB_18_, we computationally mapped out the pores with CAVER [58] for each of the FastRelaxed [59–61] predicted structures. The bottleneck radius for the predicted 16-stranded PorB_18_ is 1.55 Å (Fig. 6E), while for the 18-stranded PorB_18_ is 3.86 Å (Fig. 6F). This is slightly larger than the bottleneck radius of 3.74 Å for PorB_WT_ (Fig. 6G).

To verify the pore size experimentally, we measured the channel conductance of PorB_WT_ and PorB_18_. The channel conductance of PorB_WT_ was measured at 4.1 nS (Fig.7A). The experimental conductance for the chimeric PorB_18_ was measured to be 4.4 nS (Fig. 7A). This is slightly more than PorB_WT_ (Fig. 7B).

**Figure 7:**
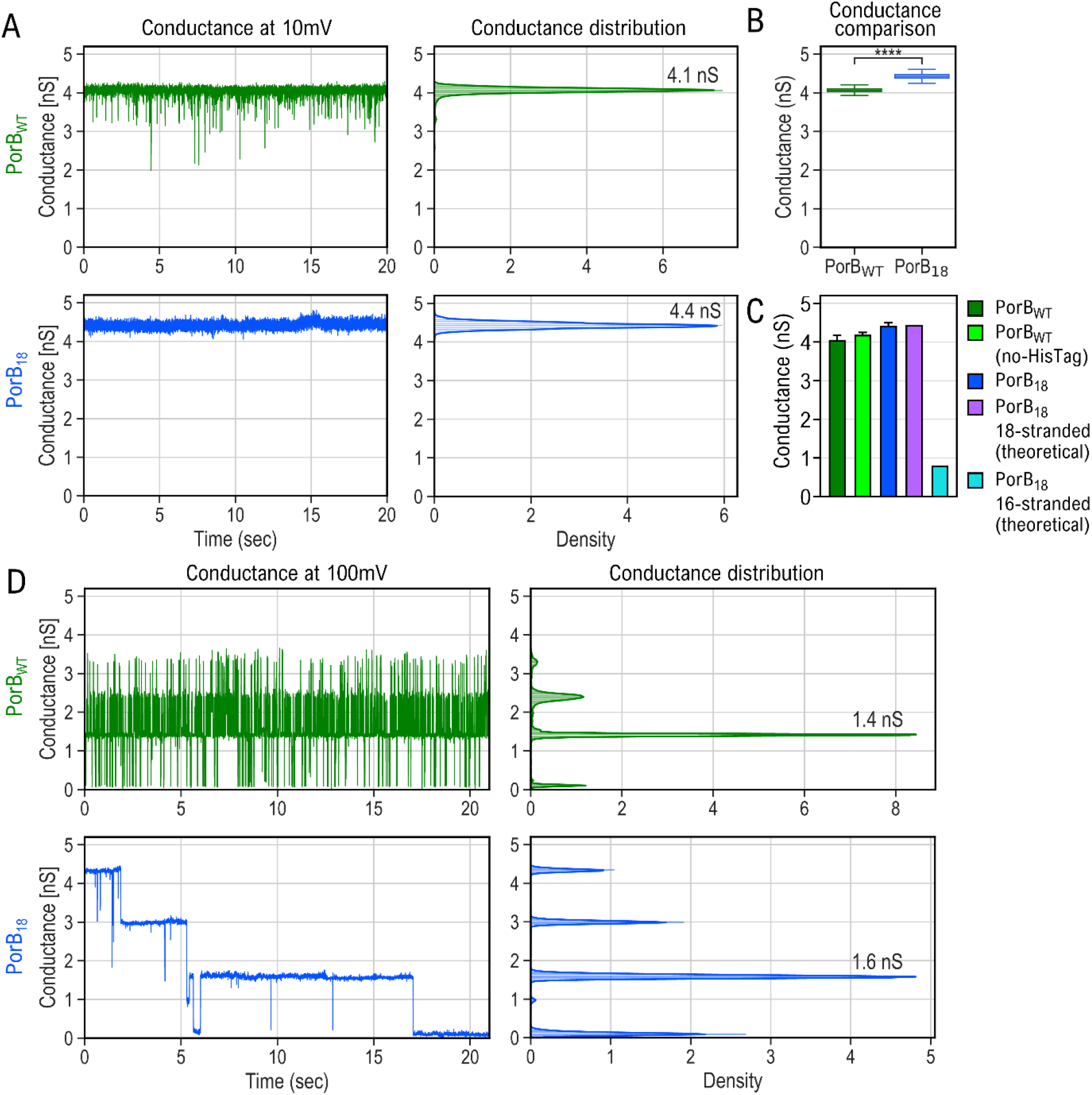
Single channel conductance of in vitro folded PorB_18_. **(A)** Single channel conductance of PorB_18_ (blue) compared to PorB_WT_ (green). The graphs on the left show the conductance over time. The graphs on the right show the conductance distribution. **(B)** Box and whisker plot comparing the conductance of PorB_WT_ (green) and PorB_18_(blue). ****: p-value <= 10^-4^ **(C)** Graph shows comparison of experimental single-channel conductance of PorB_WT_ with His-Tag (green) and without His-tag (light green), PorB_18_ (blue), theoretical shingle-channel conductance of AlphaFold 18-stranded model (magenta) and 16-stranded model (cyan) of PorB_18_. **(D)** Channel conductance at 100 mV shows channel closing for individual subunits of PorB_WT_ (green) and PorB_18_ (blue).

The conductance of PorB_WT_ shows frequent closing and opening of its channel when compared to PorB_18_ (Fig. 7A). This frequent channel closure is likely due to the presence of the N-terminal His-tag, which in PorB_WT_ leads to brief obstructions in the pore opening. We confirmed the absence of channel closure in PorB_WT_ when the N-terminal His-tag was absent (Fig. S3). The absence of channel closure in PorB_18_, even with a His-tag present, possibly indicates that either the opening of the channel for PorB_18_ is wide enough to not be affected by the His-tag or that the His-Tag is in a conformation that does not interact with the channel. Despite the artifactual noisiness of the His-tagged PorB_WT_, we compare His-tagged constructs as we could not purify PorB_18_ without the His-tag.

The theoretical conductance for PorB_18_ AlphaFold models were calculated using equation 1 and by solving the unknown variables by using the experimental conductance value of 4.2 nS for PorB_WT_ without the His-Tag (Fig. 7C) and its calculated pore size of 3.74 Å (Fig, 6G). The 18-stranded model of PorB_18_ has a theoretical conductance of 4.35 nS whereas the 16-stranded model of PorB_18_ has a theoretical conductance of 0.79 nS due to a narrower pore radius of 1.55 Å (Fig. 6E). The experimental conductance of PorB_18_ is much closer to the theoretical conductance of the 18-stranded AlphaFold model and much larger than that of the 16-stranded model (Fig. 7C). This shows that the inserted region is not only obstructing the pore but is also likely integrating into the barrel to form a larger pore size.

To rule out the possibility that the higher conductance of PorB_18_ is not from multiple PorB_18_ trimers but rather from a single trimeric PorB_18_, we measured the conductance at a higher voltage. The high voltage causes the pores of the porins to close and open at a much higher frequency. This causes multiple channels to close simultaneously, resulting in different levels of conductance. The distribution of conductance can provide insight into the number of individual porins present [34]. Both PorB_WT_ and PorB_18_ showed four levels of conductance providing evidence that the observed conductance is from a trimer (Fig. 7D). The highest level of conductance for PorB_WT_ is seen to be lower than 4.1nS because the frequency with which all three channels are open at the same time is very low, shifting the distribution of the conductance downward.

### Shortened loop constructs collapse

To evaluate the function of the long loops native to PorB and MOMP, shorter loop constructs for PorB_WT_ and PorB_18_ were created with their long loops shortened (loop 3 for PorB_WT_ and the MOMP loop 4 for PorB_18_) (Fig. 8A). We called these shorter loop constructs, PorB_SL_ and PorB_18SL._ We produced these in *E. coli* with an outer membrane targeting signal sequence. Western blot of the whole cells showed that PorB_SL_ does not express well indicating that loop L3 might be required for stable expression of PorB_WT_ (Fig. 8B). Expression and folding into micelles showed PorB_SL_ to be dimer rather than trimer (Fig. 8C). The shortened loop may be disrupting the interaction between loop L1 and loop L4 at the barrel-barrel interface (Fig. 8D).

**Figure 8:**
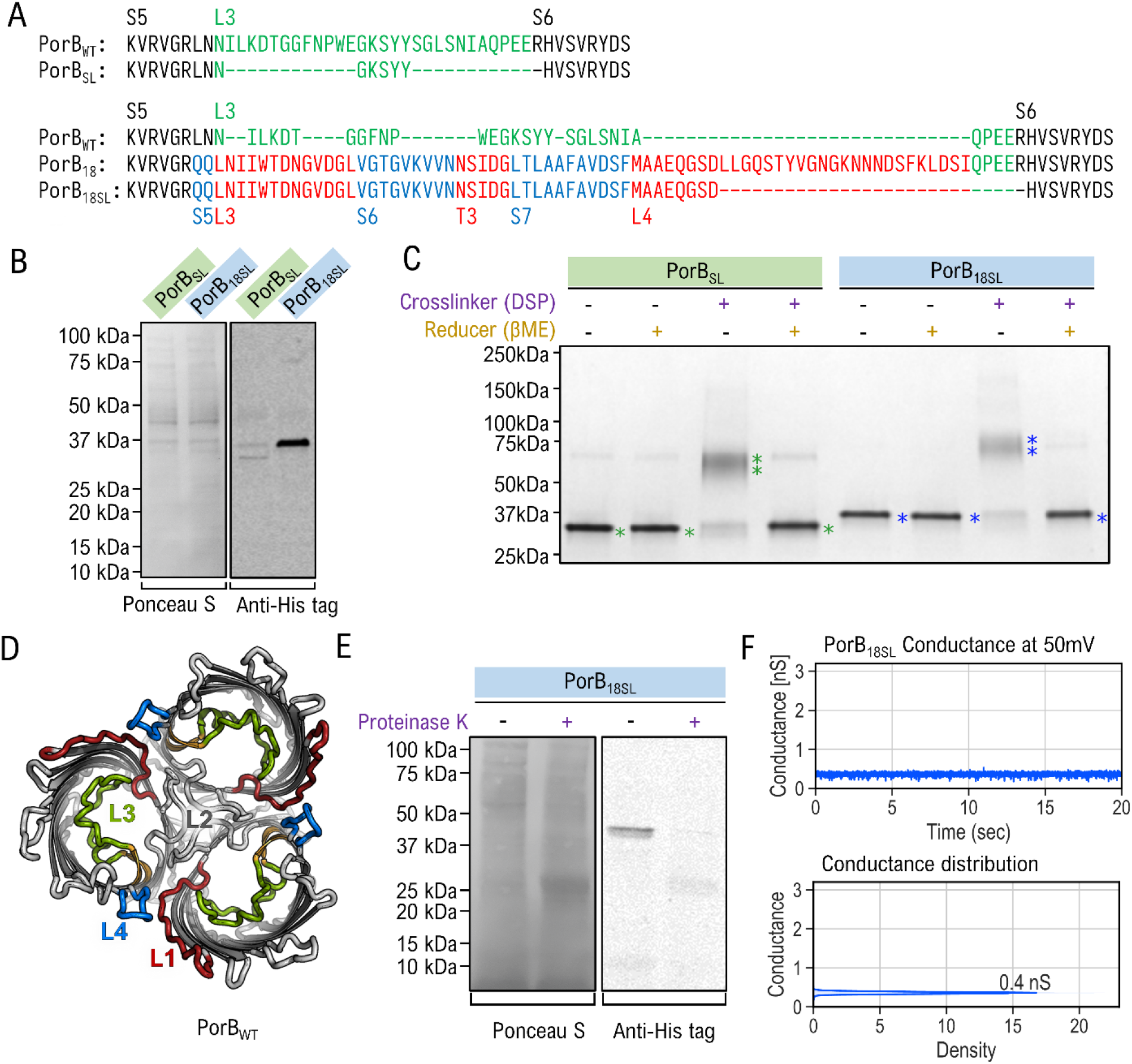
Loop shortening in PorB_WT_ and Por_18_ leads to pore collapse. **(A)** Top sequence alignment shows the short loop construct of PorB_WT_ with L3 deletion named PorB_SL_, compared with PorB_WT_. The bottom three sequences compare the short loop construct of PorB_18_ with L4 deletion named PorB_18SL_, compared with PorB_WT_. and PorB_18_. Strand sequences from PorB_WT_ are in black and loops are in green. Strand and loop sequences taken from MOMP are in blue and red respectively. **(B)** Western blot showing expression levels of PorB_SL_ and PorB_18SL_ in whole cells. **(C)** DSP crosslinking of the peak fractions from SEC. Monomeric bands are marked with (⁎) and dimers with (⁑). **(D)** Cartoon of PorB_WT_ trimer showing loops L1 to L4 (PDB ID 4aui) [39]. **(E)** Western blot showing PorB_18_ is susceptible to proteinase K digestion. **(F)** Single channel conductance of PorB_18SL_

While PorB_18SL_ expressed and folded into the *E. coli* outer membrane, it was found to be susceptible to proteinase K (Fig. 8E). Single channel conductance of PorB_18SL_ inserted into lipid bilayer membrane showed that its conductance is an order of magnitude smaller than PorB_18_ (Fig. 8F). PorB_18SL_ is likely collapsing without the long loop. Barrel collapse has been observed in simulations of engineered 16-stranded β-barrel without a lumen-lining long loop [62].

Proteinase K digestion PorB_SL_ and PorB_18SL_ folded into micelles showed more susceptibility compared with PorB_18_ (Fig. S4A, S4B, and 4B), where the short-loop mutants start to show degradation within 30 sec of digestion and are almost fully degraded after 15 minutes of digestion. MALDI-TOF of the digested proteins show two major cleavage products for both the short loop constructs with mass differences of 4.8 kDa and 9.1 kDa from the full-length proteins (Fig. S4C and D). These mass differences are similar to PorB_18_ cleavage products (Fig. 4F, peak at 35,649.5 and 31,292.8 m/z), indicating cleavage at the same site as PorB_18_. Due to the short-loop variant’s inability to form trimers (Fig. 7C), these sites are likely to be exposed. Compared to PorB_SL_, the PorB_18SL_ digestion revealed four additional cleavage products after 15 minutes, likely due to exposed proteinase K sites present in the inserted region.

CD spectra for the short loop constructs show a primarily β-sheet structure (Fig. S5A). The thermal stability of the short loop construct was comparable to PorB_WT_ measured through CD absorption at 218 nm (Fig. S5B). Interestingly, the small rise in CD absorption at 218 nm seen between 20 °C and 60 °C for PorB_WT_ (Fig. 3D) is absent in PorB_SL_ (Fig. S5B). This indicates that loop L3 undergoes temperature-dependent structural change. Secondary structure estimates from the CD spectra indicate a higher β-strand percentage for both PorB_SL_ and PorB_18SL_ (Fig. S5C) compared to the PorB_WT_ (Fig. 3E), as expected due to the removal of the loops.

## Discussion

Here, we engineer a larger pored β-barrel from PorB by inserting domains from another β-barrel protein from a different species (MOMP). Despite having a large section modified the chimeric protein PorB_18_ is found to be stable, likely due to the evolutionary relation between PorB and MOMP [21]. Outer membrane β-barrels have been engineered into larger-sized barrels through duplication of existing strands [19, 29–32]. These methods mimic the gene duplication of topology accretion in repeat proteins which are generally considered to be the predominant way of protein evolution. OMP evolution with a loop-to-hairpin mode of strand accretion had not been explored experimentally before. Our results with PorB_18_ support that this mode of strand accretion can also be used to engineer larger barrels and was likely to have been used by evolution.

Successful expression and localization of the chimeric protein PorB_18_ in the outer membrane demonstrate that the inserted region does not affect its ability to fold in bacterial membranes. The chimeric protein is also resistant to proteinase K in bacterial membranes but is susceptible to proteinase K when refolded from solubilized inclusion bodies in detergent micelles. On this basis, we can speculate that the extracellular loops are shielded by LPS in bacterial membranes but become exposed in detergent micelles. Although increased susceptibility when adding a new part to a protein sometimes indicates that that part is unfolded, the four cleavage products that are a result of PorB_18_ proteinase K digestion are not consistent with cleavage in the β-strands added to PorB_18_. Moreover, the substantial agreement between the 18-stranded AlphaFold2 predicted structure and our experimental data, including β-strand content from circular dichroism and pore size from experimental channel conductance indicate the likely incorporation of the β-hairpin into the barrel.

Biophysical assays indicate that the chimeric PorB_18_ has more β-strand content, and the strands are integrating into the β-barrel resulting in a larger pore size. While we did not have an experimental structure for the chimeric protein PorB_18_, the increase in single-channel conductance when compared to PorB_WT_ closely matches the increase in β-structure and increase in predicted pore size for the 18-stranded AlphaFold prediction for PorB_18_. This supports that the chimeric protein is 18-stranded compared to the 16-stranded wild type, enlarging the β-barrel.

The lumen-lining long loop of PorB is necessary for the stability of the protein. Deletion of loop L3 prevented PorB from trimerizing. We hypothesize that the truncated loop is preventing interaction between loop L2 and L4 of neighboring monomers, destabilizing the trimer. This can be further explored with mutational studies to verify the importance of this interaction for the trimerization of PorB_WT_.

In addition to the L2-L4 interactions, the lack of trimerization in the short loop constructs, PorB_SL_ and PorB_18SL_, could be a result of the β-barrel collapsing on itself without the support of the long loop lining the lumen of the barrel. A flattened β-barrel would only be able to interact with a single protomer and form a dimer. The small increase in conductance of a single channel in PorB_18_ (4.2 nS to 4.4 nS) and the corresponding AlphaFold structure indicate that the inserted loop region is forming a constricting loop within the lumen of the barrel. While we had expected that the deletion of this loop would result in a significant increase in conductance due to the increased barrel size and the absence of the constricting loop, we instead saw a tenfold decrease in channel conductance (0.4 nS) for PorB_18SL_. This strongly supports that the PorB_18SL_ barrel is collapsing. This barrel collapse is also the most likely explanation for PorB_SL_, where we did not see any conductivity in repeated experiments.

The fact that an extracellular loop can easily be converted into a transmembrane β-hairpin brings OMP folding questions to the fore. Future studies may clarify if the barrel folds synchronously or if the original 16 strands fold first and then the newer strands are added. Moreover, it remains possible that some loops are transiently hairpins, or some hairpins are transiently loops. Such a transition within a protein has been found within soluble proteins [63] though has not yet been found in membrane proteins. In membrane β-barrels, such a transition would bring previously unknown plasticity to the barrel radius allowing for the passage of larger solutes or acting as a gating mechanism. Finally, while we tested the loop-to-β-hairpin evolution between porins PorB and MOMP, there are other evolutionarily related pairs that could also be tested to determine how widely this method can be used.

## Conclusion

Here we show the first experimental evidence of the evolutionarily unusual topology accretion through the loop-to-hairpin transition in OMBB evolution using a PorB chimeric protein. This provides proof for one such protein evolutionary pair and opens up the avenue to explore more of such relationships in outer membrane β barrels as well as other protein structures.

## Materials and Methods

### Cloning and expression in the bacterial membrane

Plasmids for recombinant protein expression were constructed using mega primer insertion PCR [64] using the plasmid Champion pET303/CT-His vector (Invitrogen). The double-stranded DNA inserts (Integrated DNA Technologies) consisted of the codon-optimized signal peptide from *E. coli* OmpA protein, a 6x histidine tag, a TEV protease cleavage site, the recombinant protein, two stop codons, and two 50-nucleotide homologous regions from the vector flanking on each side. NEB Turbo *E. coli* strain (New England Biolabs) was used for cloning and plasmid propagation.

*E. coli* ΔDegP strain (KS476 strain acquired from CGSC)[65] was modified by integration of λDE3 prophage using the Novagen λDE3 Lysogenization Kit (MilliporeSigma) to allow for the use of T7 polymerase-based plasmid vectors. The plasmid vector for the recombinant protein was transformed on LB agar plates with carbenicillin as the selection antibiotic. A single colony was resuspended in 2xYT media with antibiotics grown overnight at 37 °C at 250 RPM shaking incubator. Protein expression was achieved in these overnight cultures through the leaky expression of the pET303 vector (without any inducing agent).

## Outer membrane fractionation

Overnight cultures were collected by centrifugation at 3000 g for 10 min at 4 °C. The cell pellets were then resuspended in lysis buffer (50 mM Tris, pH 7.4, 200 mM NaCl, 10 mM MgCl_2_, 5 mM benzamidine, 0.5 mM phenylmethylsulfonyl fluoride (PMSF), 5 μg/mL DNase). Resuspended cells were freeze-thawed thrice by freezing them at -80 °C and rapidly thawing at 37 °C. Resuspended cells were then lysed using a sonicator (Qsonica Q500 Sonicator) submerged in an ice bath. Sonication was carried out at 40% amplitude with a pulse of 2 sec on and 4 sec off for a total ‘on time’ of 5 min. The outer membrane was fractionated as previously described [66] with some modifications. Specifically, the inner membrane solubilization buffer was modified to 20 mM sodium phosphate, pH 7.4, 5 mM benzamidine, and 2% *N*-lauroylsarcosine. The outer membrane solubilization buffer was modified to 20 mM NaPi pH 6.5, 100 mM NaCl, 5 mM benzamidine, 2 µM lysozyme, and 1% LDAO. Benzamidine was required in all buffers for the inhibition of the endogenous protease OmpT present in the ΔDegP (DE3) strain. The outer membrane was solubilized for 30 min at room temperature and then separated from non-soluble fractions by centrifugation at 15,600 g for 30 min. Fractions were analyzed by Western blot.

## Proteinase K assay in permeabilized bacterial membrane

The proteinase K assay was conducted as previously described [50] with some modifications. In brief, 5 mL of overnight cultures expressing the recombinant protein were collected by centrifugation (3000 g, 5 min, 4 °C). The cells were permeabilized by resuspending the cell pellets in the permeabilization buffer (33 mM Tris pH 8.0, 40% sucrose, 2 μM lysozyme, and 2 mM EDTA) and incubating on ice for 20 min. Half of the cell suspensions were treated with 3.5 μM proteinase K (Sigma-Aldrich) and half were left untreated as a control. The digestion was stopped with 1 mM PMSF after 20 min. The cell suspensions were precipitated with 10% TCA. The precipitates were collected by centrifugation at 20,000 g for 5 min. The precipitates were washed once with 100% acetone and twice with 50 mM Tris pH 8.0, and 200 mM NaCl. Precipitates were mixed with 2x Laemmli sample buffer (BioRad) and boiled for 10 min at 98 °C to prepare them for western blot.

## SDS-PAGE and Western blot

Purified proteins, cell fractions, or whole cells were mixed with 2x Laemmli sample buffer (BioRad) and loaded onto a 4-20% polyacrylamide gel (Bio-Rad). Electrophoresis was carried out at 300 V, for 15 min. Gels were stained with ReadyBlue Coomassie protein gel stain (MilliporeSigma) and imaged on Gel Doc EZ imaging system (BioRad)

For the Western blot, after the electrophoresis step, gels were sandwiched with 0.2 μm nitrocellulose blotting membrane (BioRad), and the transfer was carried out at 100 V for 30 min on ice. The transfer buffer was prepared with 10x Tris-glycine buffer (BioRad) and 10% ethanol. To verify the transfer, blots were stained with 0.2% (w/v) Ponceau S in 5% glacial acetic acid.

Blots were washed thrice in TBST (20 mM Tris, pH 7.5, 150 mM NaCl, 0.1% Tween 20) and blocked with TBST with 1% gelatin from cold water fish skin (MilliporeSigma). The following antibodies were diluted as per manufacturer recommendation with TBST with 1% gelatin: THE Anti-His tag monoclonal mouse antibody (GenScript), anti-*E. coli* SurA polyclonal rabbit antibody (Cusabio), IRDye 800CW goat anti-mouse IgG secondary antibody (LI-COR Biosciences), IRDye 680RD donkey anti-rabbit IgG secondary antibody (LI-COR Biosciences). anti-*E. coli* TolC was purified from rabbit sera and diluted to 1:2500 with 1% gelatin in TBST. All incubations were carried out at 37 °C for 30 min, followed by three 10 min rinses with TBST at 37 °C. Blots were imaged with LI-COR Odyssey infrared imager at 700 nm and 800 nm.

## Protein expression, refolding, and purification

Recombinant proteins were expressed in *E. coli* BL21(DE3) as inclusion bodies by removing the OmpA signal sequence. Protein expression was induced at a culture density of 0.6 OD600 with 1 mM IPTG. Cells were collected by centrifugation at 4000 g for 30 min. Inclusion bodies were harvested by lysing the cell suspension in ice-cold lysis buffer (50 mM Tris pH 8, 200 mM NaCl, 1 mM EDTA, 1 mM PMSF) with 2% TritonX-100. Lysis was carried out by sonication at 50% amplitude, with a pulse of 2 sec on and 2 sec off for a total ‘on time’ of 5 min per liter of cells, submerged in an ice bath. Cells were washed thrice by centrifugation at 11000 g for 10 min, followed by resuspension in ice-cold lysis buffer without TritonX-100.

Inclusion bodies of his-tagged protein were solubilized in binding buffer (20 mM NaPi, pH 6.5, 100 mM NaCl, 20 mM imidazole, 7.2 M urea). Solubilized inclusion bodies were purified by metal affinity chromatography using a precharged Ni Sepharose 6 Fast Flow prepacked column (Cytiva) on an ÄKTA pure FPLC system (Cytiva). Protein was eluted with a gradient of 0 – 50 % elution buffer (20 mM NaPi, pH 6.5, 100 mM NaCl, 500 mM imidazole, 7.2 M urea). Protein eluent was concentrated using 10 kDa Amicon Ultra-15 centrifugal filter units (MilliporeSigma) to a maximum concentration of 20 mg/mL for PorB_WT_ and 7 mg/mL for the mutant proteins. The protein concentrate was refolded by mixing it with refolding buffer (10% LDAO, 20 mM NaPi pH 6.5, 100 mM NaCl, 1 mM EDTA) at a 1:2 ratio [52]. Freshly folded protein was separated from aggregates using size exclusion chromatography using a prepacked Superdex 200 10/300 GL column (Cytiva) equilibrated with running buffer (0.1% LDAO, 20 mM NaPi pH 6.5, 100 mM NaCl, 1 mM EDTA).

## Crosslinking assay

Protein eluates from SEC were crosslinked at 2.5 µM concentration with 250 µM DSP (Thermo Scientific) at room temperature for 30 min. The reactions were quenched with 50 mM final concentration of Tris-HCl pH 7.5 for 15 min. Aliquot from each reaction was mixed with 2x Laemmli SDS sample buffer (Bio-Rad) for SDS-PAGE analysis.

## Circular dichroism

Samples for circular dichroism (CD) were prepared by buffer exchanging the purified protein into CD buffer (0.1% LDAO, 20 mM NaPi pH 6.5, 100 mM NaF) using a Superdex 200 10/300 GL column (Cytiva). The eluent was concentrated using 30 kDa Amicon Ultra-0.5 centrifugal filter devices (MilliporeSigma) to a concentration of 1 mg/mL. Protein samples were loaded onto a 0.1 mm pathlength Suprasil quartz cuvette (Hellma) for CD measurement. CD spectra were collected using a JASCO J-815-150S CD spectrometer with the following parameters: wavelength range of 260-175 nm, speed of 100 nm/min, data pitch of 0.5 nm, D.I.T of 2 sec, bandwidth of 1 nm, and 7 accumulations for each sample. The secondary structure was estimated using JASCO’s multivariate secondary structure estimation program. Thermal melt curves were generated by using the spectrophotometer with a JASCO PTC-423 Peltier controller accessory. CD spectra were collected at different temperatures starting from 20 °C and up to 90 °C at 5 °C increments.

## Single channel conductance

The single-channel setup includes a BC-535 bilayer clamp amplifier from Warner Instruments, National Instruments’ PCI-6221 DAQ card for data acquisition, and National Instruments’ BNC 2110 digitizer. Data was recorded using WinEDR software (J. Dempster, University of Strathclyde, UK). The bilayer chambers consist of Warner instruments BCH-M13 derlin chamber and CD13A-200 derlin cuvette with 200 µm aperture. 1.4 mL 1 M KCl was added to each chamber as well as the electrode chambers. The electrode chambers were connected to the main chambers using 3 M KCl, 2% agar bridges. The *cis*-chamber was connected to the input and a *trans*-chamber was connected to the ground. The bilayer was formed by ‘painting’ a solution of 10 mg/mL 1,2-diphytanoyl-sn-glycero-3-phosphocoline in *n*-decane over the aperture. Bilayer formation was confirmed by observing an increase in membrane capacitance to 80 pF – 110 pF. 2 µL of 1 mg/mL protein sample was added to the *cis*-chamber near the aperture. A voltage of 100 mV was applied for incorporating the protein into the bilayer. The theoretical conductance was calculated using the following previously derived equation [67]:

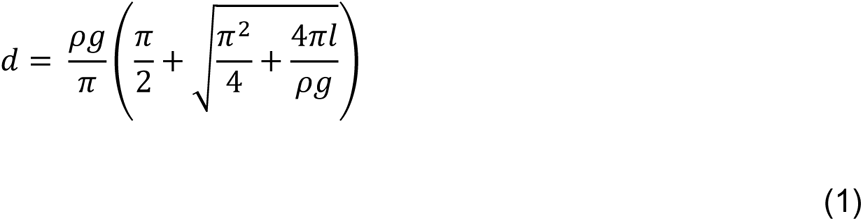

Where, *d* is the diameter of the pore, *g* is the conductance of the pore, *ρ* is the resistivity of the buffer, and *l* is the length of the pore.

## Proteinase K assay in micelles and mass spectrometry

For proteinase K digestion in micelles, purified proteins at a concentration of 2 μM were treated with 0.1 μM of proteinase K and placed on ice. PMSF was added at 1 mM concentration to stop the digestion at specific time points. Samples were mixed with 2x Laemmli sample buffer (BioRad) for SDS-PAGE analysis.

Proteinase K treated samples were analyzed using matrix-assisted laser desorption/ionization-time of flight (MALDI-TOF) mass spectrometry. Samples were made acidic by adding TFA to 0.1% final concentration and then desalted using a ZipTip with C_4_ resin (MilliporeSigma). The desalted protein was step-wise eluted with solvents with varying acetonitrile concentrations. First, solvent TA20 (20:80 [v/v] acetonitrile/0.1% TFA in water) was used to elute smaller mass peptides, followed by solvent TA30 (30:70 [v/v] acetonitrile/0.1% TFA in water) to elute smaller mass proteins, and finally with TA50 (50:50 [v/v] acetonitrile/0.1% TFA in water).to elute larger mass proteins. The eluent was mixed with α-cyano-4-hydroxycinnamic acid (HCCA) matrix solution (supersaturated HCCA solution in solvent TA30) at a 1:1 ratio. Samples were prepared by seed-layer method [68]. First, 1 mg/mL HCCA in 100% acetonitrile was deposited on a 96-spot, polished, stainless steel target plates (Bruker Daltonics) and dried. Then 1 µL of each protein matrix mixture was deposited onto the dried HCCA spots. For calibration, Protein Standard II (Bruker Daltonics) was prepared with the HCCA matrix solution and 1 µL was deposited onto a dried HCCA spot, next to the protein sample spots. The protein mass to charge spectra were acquired on a MicroFlex LT MALDI-TOF system (Bruker Daltonics) with 1000-2000 shots using the instrument’s 337 nm nitrogen laser.

## Structure prediction and pore mapping

To predict the structure of PorB_18_, we used AlphaFold v2.3.1 [55, 56] through ColabFold v1.5.2 [54], using MOMP as a template (See supplementary text for configuration). The top two predictions with the highest pLDDT were relaxed in RosettaScripts [61] FastRelax protocol [59, 60]. The relaxed structure with the lowest Rosetta score was used for selecting for mapping its pore using the CAVER 3.03 PyMol plugin [58]. The starting point for the tunnel was selected on the periplasmic side of the protein. The correct path was selected visually for the bottleneck radius of the pore.

## Data analysis and graphs

Single-channel data analysis and proteinase K digest product calculations were done using Python 3.10.7 and the Pandas 1.3.4 library [69]. Graphs for these were generated using Python libraries Seaborn v0.11.2 [70] and Matplotlib v3.5.1 library [71]. Chromatography and MALDI-TOF data were graphed using GraphPad Prism 9.5.1 (GraphPad Software, San Diego, California USA, www.graphpad.com).

## Acknowledgments

We thank Ayotunde Paul Ikujuni, Jimmy Budiardjo, Ryan Feehan, and Daniel Montezano for their helpful insights and discussion. We also gratefully acknowledge NIGMS awards DP2GM128201, R01GM148583, P20GM113117, and P20GM103418 and NSF 2226804 to JSGS.

## Abbreviations

OMP: Outer membrane proteins
MOMP: Major outer membrane protein
SDS-PAGE: Sodium dodecyl sulfate-polyacrylamide gel electrophoresis
CD: circular dichroism
SEC: size exclusion chromatography
MALDI-TOF: matrix-assisted laser desorption/ionization-time of flight
pLDDT: predicted local distance difference test
PMSF: phenylmethylsulfonyl fluoride
PDBID: protein data bank identification
PK: proteinase K

