## Supplemental Figures for "Evolutionary engineering a larger porin using a loop-to-hairpin mechanism"

### Evolutionary Engineering of a larger Pore based on a loop to hairpin mechanism

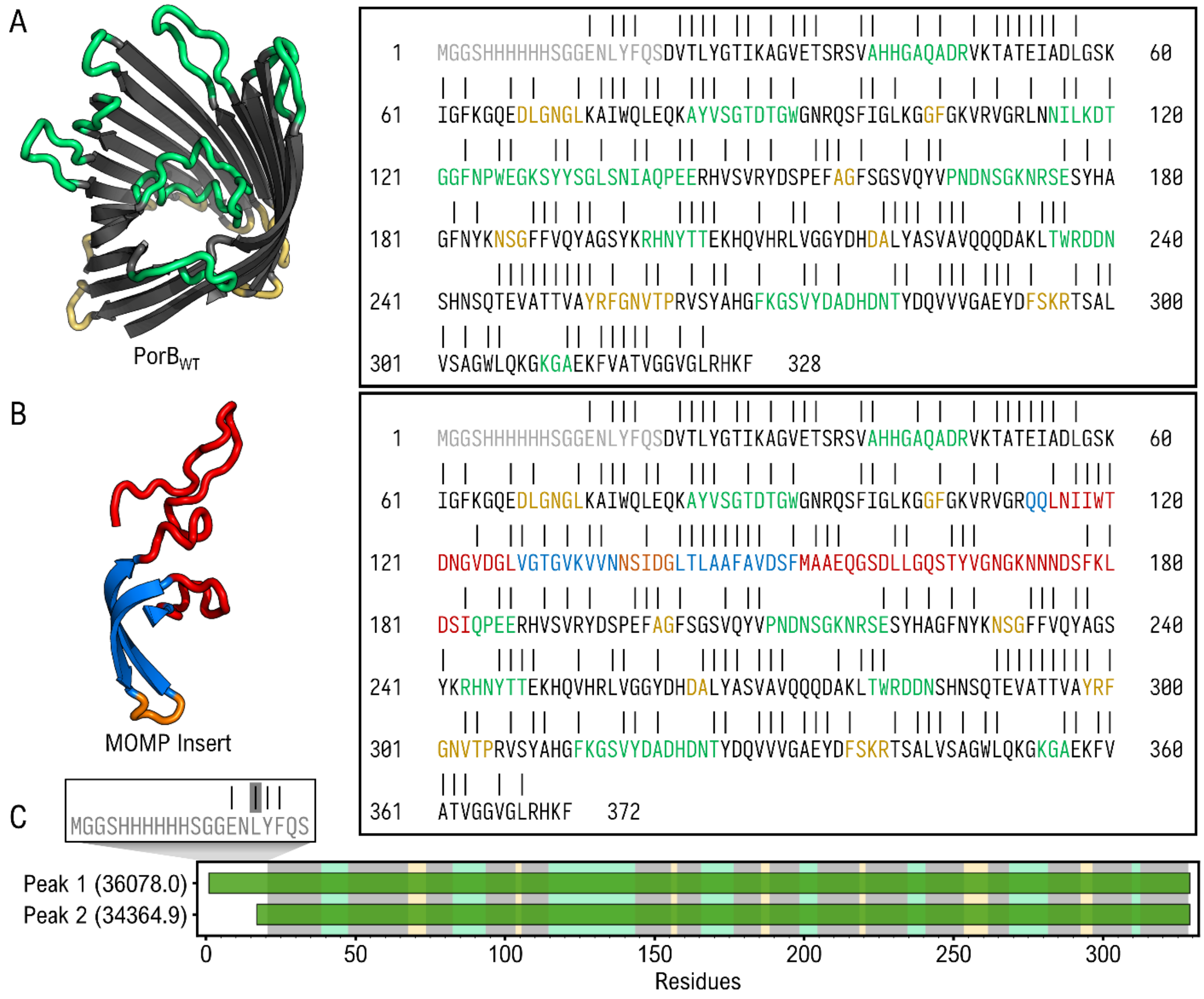


**Figure S1: Predicted Proteinase K cleavage sites of PorB$_{WT}$ and PorB$_{18}$:** Cleavage sites are predicted using PeptideCutter from ExPASy Server [1] for **(A)** PorB$_{WT}$ and **(B)** PorB$_{18}$. Vertical lines indicate where Proteinase K can potentially cut the peptide bond at the C-terminal end of the amino acid. Amino acid colors represent both the secondary structure and the original protein used to create the chimeric protein: Disordered peptide tag (gray); Beta-strand s from PorB (black), extracellular loops from PorB (green), and turns from PorB (yellow); Beta-strands from MOMP (blue), extracellular loops from MOMP (red), and periplasmic turns from MOMP (orange). **(C)** Graph showing sequence range for PorB$_{WT}$ MALDI-TOF peaks from Fig. 4E. The x-axis shows the residue numbers. The outer box shows the amino acid sequence containing the cleavage site highlighted in grey.

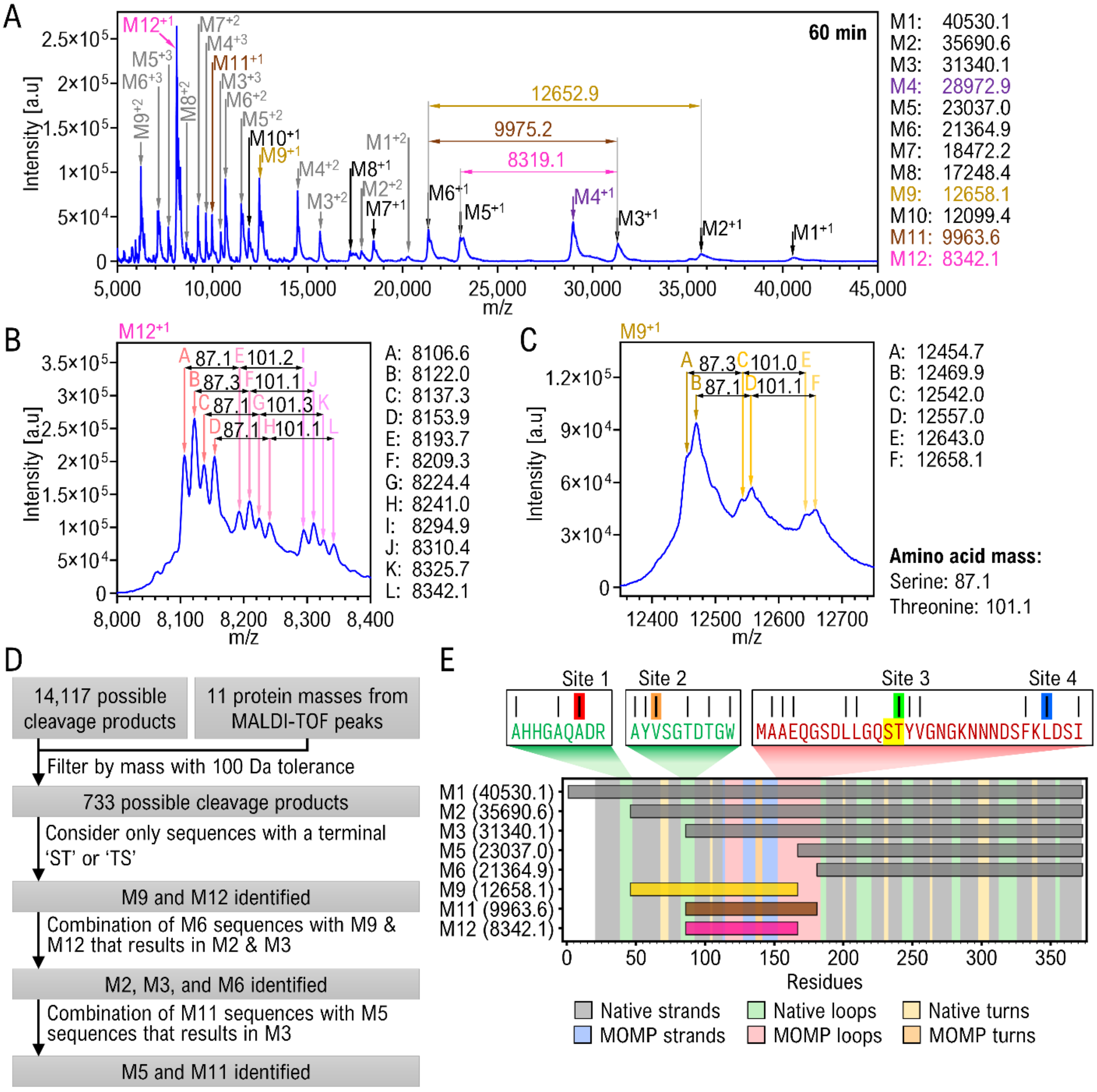


**Figure S2: Identification of MALDI-TOF peaks of Proteinase K digest of PorB$_{18}$. (A)** Extended MALDI-TOF spectra of the 60 min proteinase K digest of PorB$_{18}$ from 50,00 to 45,000 mass by charge ratio (m/z) (this includes the 20,000 to 45,000 shown in Fig 6). Unique protein masses are listed on the right side of the graph. Peaks are identified by arrows and their corresponding protein masses are shown above the arrows. We manually assigned multiply charged m/z species and found that of the 25 peaks, only 12 are a result of a distinct mass. Charge for each protein mass is shown as superscript. Proteinase K mass is shown in purple and multiple charged masses are shown in grey. Mass differences between selected peaks are shown with horizontal double arrows and are colored magenta, brown, and yellow. Protein masses associated with these mass differences are colored with the same color. This analysis allowed us to conclude that M2 is cleaved into M6 and M9, and M3 is cleaved into both M6 and M11 as well as M5 and M12, Zoomed in MALDI-TOF spectra shows peaks associated with mass **(B)** M12 and **(C)** M9. Sub-cleavage peak masses are listed next to each of the graphs showing a pervasive difference of the mass of threonine and serine. **(D)** Flow chart of our process for sequence identification of protein masses from MALDI-TOF peaks. **(E)** Graph showing the sequence range of each of the identified protein masses from MALDI-TOF peaks, represented with horizontal bars. The x-axis shows the residue number and the left and right edges of the bars represent the N- and C-terminal residues. The colored vertical bars represent the secondary structure for the residue at that position. The outer box shows the amino acid sequence containing the 'ST' sequence highlighted in yellow. The horizontal lines above the sequences mark the possible cleavage sites. The vertical lines above the sequences highlighted in red, orange, green, and blue represent cleavage sites 1, 2, 3, and 4 shown in Fig. 6.

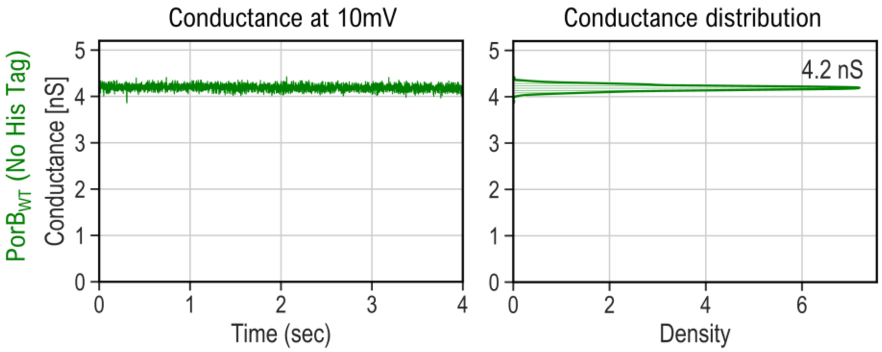


**Figure S3: Single channel conductance of PorB$_{WT}$ without HisTag.** The graph on the left shows the conductance over time. The graph on the right shows the conductance distribution.

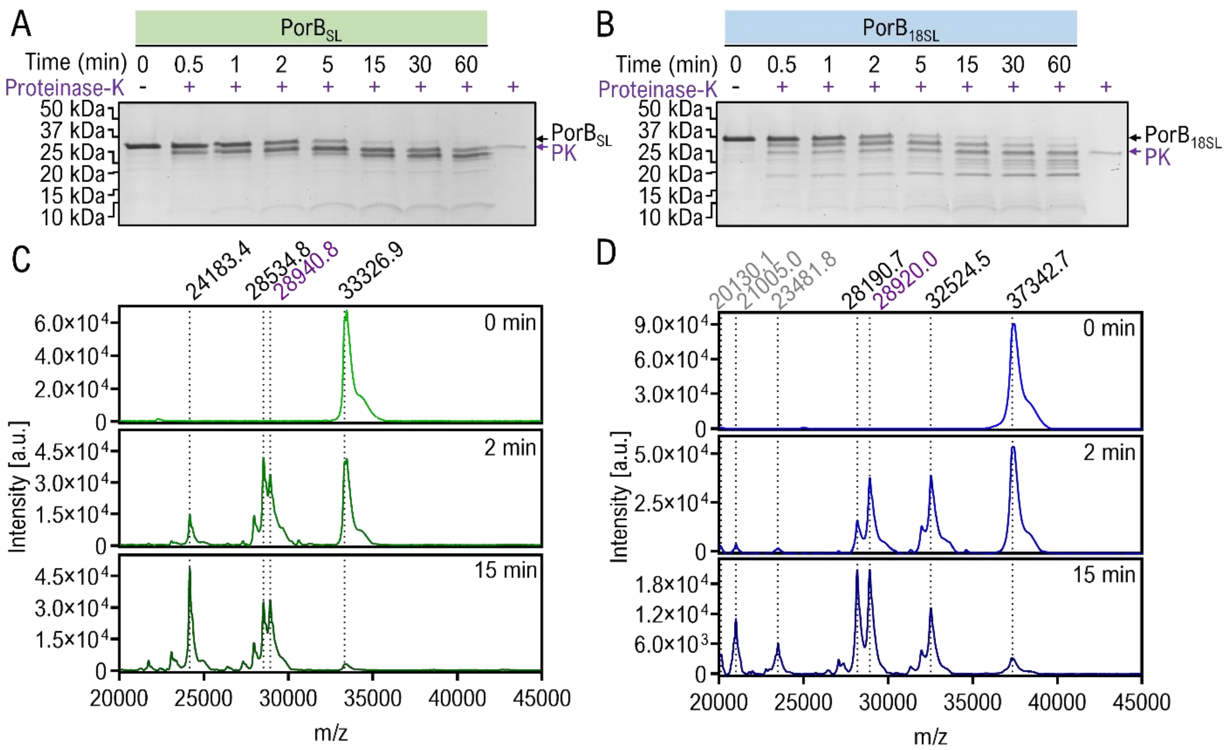


**Figure S4: Proteinase K digestion of in vitro folded PorB$_{SL}$ and PorB$_{18SL}$.** SDS-PAGE of in vitro folded proteins **(A)** PorB$_{SL}$ and **(B)** PorB$_{18SL}$, treated with Proteinase K for different durations. MALDI-TOF of **(C)** PorB$_{SL}$ (green) and **(D)** PorB$_{18SL}$ (blue) treated with Proteinase K for different durations. The numbers above the graph are the mass-by-charge ratio of the peaks. Peaks indicated by purple numbers are Proteinase K which has a molecular weight of 28.9 kDa. Peaks indicated by grey numbers are peaks that appear after 15 minutes of digestion of PorB$_{18SL}$.

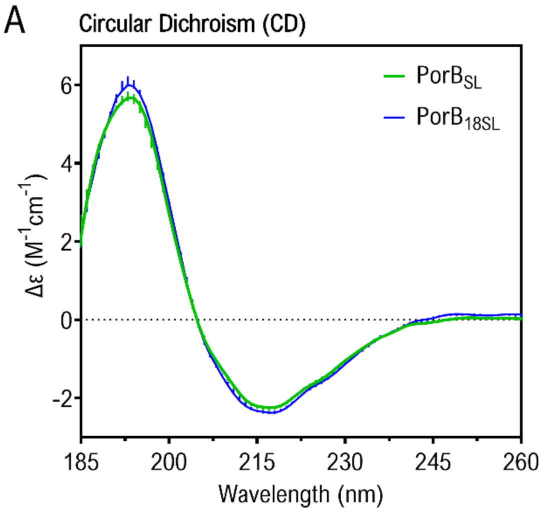


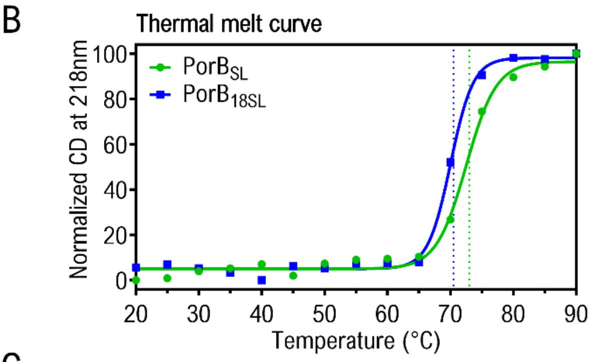


C

Secondary structure estimation

| | α-helix | β-sheet | Turns | Others |
|---|---|---|---|---|
| $PorB_{SL}$ | 7.3 ± 4.8 | 61.8 ± 3.5 | 8.8 ± 1.3 | 22.2 ± 0.9 |
| $PorB_{18SL}$ | 3.2 ± 2.9 | 67.3 ± 1.5 | 9.2 ± 1.8 | 20.4 ± 2.9 |

**Figure S5: Crosslinking and circular dichroism of $PorB_{SL}$ and $PorB_{18SL}$: (A)** CD spectra of the proteins with error bars indicated by vertical lines. **(B)** Normalized CD spectra at 218 nm showing thermal unfolding of $PorB_{SL}$ (green) and $PorB_{18SL}$ (blue). Line represents the regression curve with sigmoidal fit. Dotted vertical lines show the Tm. **(C)** Secondary structure estimation from the CD spectra.

**Computational codes:**

ColabFold Configuration:


```
{
    "num_queries": 1,
    "use_templates": true,
    "num_relax": 0,
    "msa_mode": "mmseqs2_uniref_env",
    "model_type": "alphafold2_multimer_v3",
    "num_models": 5,
    "num_recycles": 0,
    "recycle_early_stop_tolerance": null,
    "num_ensemble": 1,
    "model_order": [
        1,
        2,
        3,
        4,
        5
    ],
    "keep_existing_results": false,
    "rank_by": "multimer",
    "max_seq": 32,
    "max_extra_seq": 64,
    "pair_mode": "unpaired",
    "host_url": "https://api.colabfold.com",
    "stop_at_score": 100.0,
    "random_seed": 0,
    "num_seeds": 4,
    "recompile_padding": 10,
    "commit": "05c0cb38d002180da3b58cdc53ea45a6b2a62d31",
    "use_dropout": false,
    "use_cluster_profile": false,
    "use_fuse": true,
    "use_bfloat16": true,
    "version": "1.5.2"
}
```
